# Targeting PI3K Signaling for Broad Inhibition of β-Coronavirus Infection

**DOI:** 10.1101/2025.06.22.660949

**Authors:** Jinchao Xing, Qian Wang, Nanjun Chen, Ashley Nutsford, Blake Sullivan-Hill, John A Taylor, Gordon Rewcastle, Jack U Flanagan, Shengjin Li, Peter R Shepherd, Jianbo Yue, Natalie E Netzler

## Abstract

The phosphatidylinositol 3-kinase (PI3K) signaling pathway plays a central role in regulating key cellular processes such as survival, metabolism, and immune responses. Aberrant activation of this pathway is associated with tumorigenesis, and several PI3K inhibitors have been developed as anticancer agents. Emerging evidence suggests that viruses, including β-coronaviruses, have evolved mechanisms to exploit host PI3K signaling for their replication and immune evasion. In this study, we evaluated the antiviral efficacy of a panel of PI3K inhibitors against β-coronaviruses, including mouse hepatitis virus (MHV), human OC43 (HuCoV-OC43) and four major SARS-CoV-2 variants using both cell line and organoid models. Our findings reveal that these compounds exhibit low micromolar potency in inhibiting viral replication. Notably, the inhibitor C20 (PWT33597) demonstrated broad-spectrum activity against multiple β-coronaviruses, including SARS-CoV-2, MHV, and HuCoV-OC43, in conventional cell lines as well as in air–liquid interface (ALI)-cultured, differentiated primary human nasal and bronchial epithelial cells. Given that cytokine storm is a major contributor to SARS-CoV-2–related multiorgan failure and mortality, we further explored the impact of PI3K inhibition on host inflammatory responses. We found that MHV infection markedly increased cytokine expression in 17CL-1 fibroblasts and RAW264.7 macrophages. Interestingly, treatment with C20 further amplified cytokine production in this context, suggesting complex immunomodulatory effects that warrant further investigation. Together, our findings support the therapeutic potential of repurposing PI3K inhibitors as broad-spectrum antivirals. These compounds not only suppress viral replication but may also influence host immune responses, providing a promising avenue for intervention against current and emerging coronavirus threats.

## 1. Introduction

Coronaviruses pose a persistent health threat to humans and other animals. The *Coronaviridae* are enveloped single-stranded, positive sense RNA viruses, comprising SARS-CoV-2 and human betacoronavirus OC43 (HuCoV-OC43), as well as other significant human pathogens including HCoV-229E and MERS [1]. Since its emergence in 2019, SARS-CoV-2 continues to cause significant morbidity and mortality worldwide, with over 7 million SARS-CoV-2 deaths recorded to date [2]. Given the heavy societal burden from SARS-CoV-2 and other coronaviruses, intense interest and resources have gone into developing tools to prevent or treat infection. Even with several approved vaccines and antivirals now available globally, the continued evolution of SARS-CoV-2 and waning host immunity have rendered vaccine protection short-lived and incomplete against symptomatic disease [3] with challenges for antiviral coverage for all populations [4].

Although HCoV-OC43 usually causes mild respiratory tract infections compared to MERS and SARS-CoV-2, it has also been associated with serious disease in older-aged adults and very young children with resulting complications including bronchitis, pneumonia and encephalitis [5, 6]. HCoV-OC43 has been implicated in several outbreaks and constantly circulates within the human population as a major etiological agent of the ‘common cold’ [7-9]. At this time there are no specific antivirals available to treat HCoV-OC43 infections and only supportive care can be offered.

As obligate parasites, coronaviruses have co-evolved with their hosts to subvert many cellular functions and pathways to support the viral lifecycle. As a central controller of cell growth, survival and metabolism, phosphoinositide 3-kinases (PI3K) pathways are amongst the most targeted by different viral families, despite diverse replication strategies [10]. Viruses interact with the PI3K pathway to support various stages in their lifecycle ranging from cell entry, immune evasion, pre-mRNA splicing, viral protein production and trafficking, through to temporal control of apoptosis [10-12]. Due to high viral genome plasticity, viruses have co-evolved to target diverse points throughout the PI3K pathway to control host cell function and promote viral replication. As the PI3K/AKT/mTOR signaling pathway is central to many diverse cellular functions from protein translation to the antiviral immune response [13-15], viruses have evolved to delicately balance control without triggering rapid cellular stress responses and viral clearance by the host immune cascade [10, 11].

PI3Ks are categorized into three classes (I-III) based on their substrates, structure, and function. Class I PI3K enzymes are well characterized and are further classified into subtypes A and B depending on their regulatory action [16]. Class IA PI3K enzymes are heterodimers comprising one catalytic subunit (p110α, p110β, p110δ) and one regulatory subunit (p85α, p85β, p55α, p55γ, p50α; reviewed in [16, 17]). Class IB PI3Ks also exist as heterodimers comprising the p110γ catalytic subunit with a regulatory subunit (p101, p87, p84) [7,8]. Small-molecule inhibitors that target PI3Ks can include pan-inhibitors, isoform-specific and dual PI3K/mTOR inhibitors [16].

The persistent activation of the PI3K/AKT/mTOR pathway has been linked to cancer and therefore has been successfully targeted for the development of anticancer therapeutics [18-21]. To date, several PI3K inhibitors have been FDA approved for chemotherapy including Alpelisib, Copanlisib, Duvelisib, Idelalisib, and Umbralisib with several others in various stages of development [16, 22].

Several studies have shown that inhibition of the PI3K/AKT/mTOR pathway impacts on viral replication [12]. A number of PI3K inhibitors possess broad-spectrum antiviral activities, inhibiting replication of diverse viruses including influenza [23, 24], SARS-CoV-2 [25], MERS [26], Ebola virus [27], herpesviruses [28], rotavirus [29], human metapneumovirus [30], West Nile virus [31], and hepatitis C virus [32] amongst others. Two known exceptions to these antiviral effects are the unrelated hepatitis B and hepatitis E viruses, which both showed elevated levels of replication with PI3K/mTOR pathway inhibition in cell culture studies [33-35].

Significant efforts have gone into the development of safe and effective antivirals to combat SARS-CoV-While we have several clinically approved antivirals available such as orally administered Molnupiravir and Nirmatrelvir (Paxlovid) and intravenously administered Remdesivir [4], these antivirals have side effects and contraindications, and all are used currently as monotherapies, which increases the risk of the virus evolving towards resistance [4]. Therefore, additional safe and effective antivirals are needed to combat the clinical burden created by SARS-CoV-2, and for use in various combinations with other antivirals to raise the barrier to resistance [36].

Drug repurposing has the advantage over traditional drug discovery in that compounds have often progressed through safety and tolerability clinical trials, significantly reducing development resources and time to market [37]. Several PI3K inhibitors previously developed against cancer have published efficacy against SARS-CoV-2 in various cell and animal models [25, 38-40]. For example, the approved anti-cancer drug Capivasertib, and the preclinical candidates Pictilisib and Omipalisib demonstrated antiviral effects against SARS-CoV-2 *in vitro* [25, 39]. Another PI3K inhibitor candidate for SARS-CoV-2 treatment is the FDA approved cancer drug Duvelisib, which was shown in a clinical trial to reduce hyperinflammation in established SARS-CoV-2 cases to ameliorate pneumonia [41], and appears to disrupt virus-host interactions between the host DNA damage responder protein PARP1 and the viral Spike protein [42]. These examples demonstrate the promise of PI3K inhibitors as potential coronavirus antivirals. Additionally, hyperactivation of PI3K/Akt/mTOR has been correlated with SARS-CoV-2 infection severity, supporting the inhibition of this pathway as potential SARS-CoV-2 therapies [43].

Despite the clinically approved antivirals for SARS-CoV-2, there are as yet no approved therapies available for other coronaviruses such as HCoV-OC43, and we urgently require more safe and effective antivirals to combat the current clinical burden. Importantly, we also require a range of broad-spectrum antivirals ready to use as a first line of defense against any novel or re-emerging coronaviruses, to allow time until other tools such as vaccines can be developed. It has been widely accepted that another coronavirus could likely cause another future pandemic [44], and therefore we need a range of broad-spectrum anti-coronavirus treatments in development, to ensure protection of life while vaccines are developed against novel coronavirus pathogens.

This study examined a number of commercial and novel PI3K inhibitors in various stages of development for antiviral activities against β-coronaviruses, including murine hepatitis virus, human coronaviruses HCoV-OC43 and SARS-CoV-2 (Wuhan, Mu, Delta, and Omicron variants) using cell line and organoid models. The aims of this study were to establish the antiviral efficacy of PI3K inhibitors as a platform for antiviral development with broad-spectrum activity against viruses from the *Coronaviridae*.

## 2. Materials and Methods

### Compounds

All test compounds and controls are outlined in Table 1. The PI3K inhibitor test compounds C2, C9 and C20-23 were produced at the Auckland Cancer Society Research Centre (Table 1). The commercial PI3K inhibitors examined against MHV were purchased from MedChemExpress, and the controls were purchased as follows: DMSO and Berbamine dihydrochloride (BBM) from Sigma Aldrich, and Remdesivir (RDV) from Med Chem Express. All structures are shown in Table 1 below.

**Table 1.**
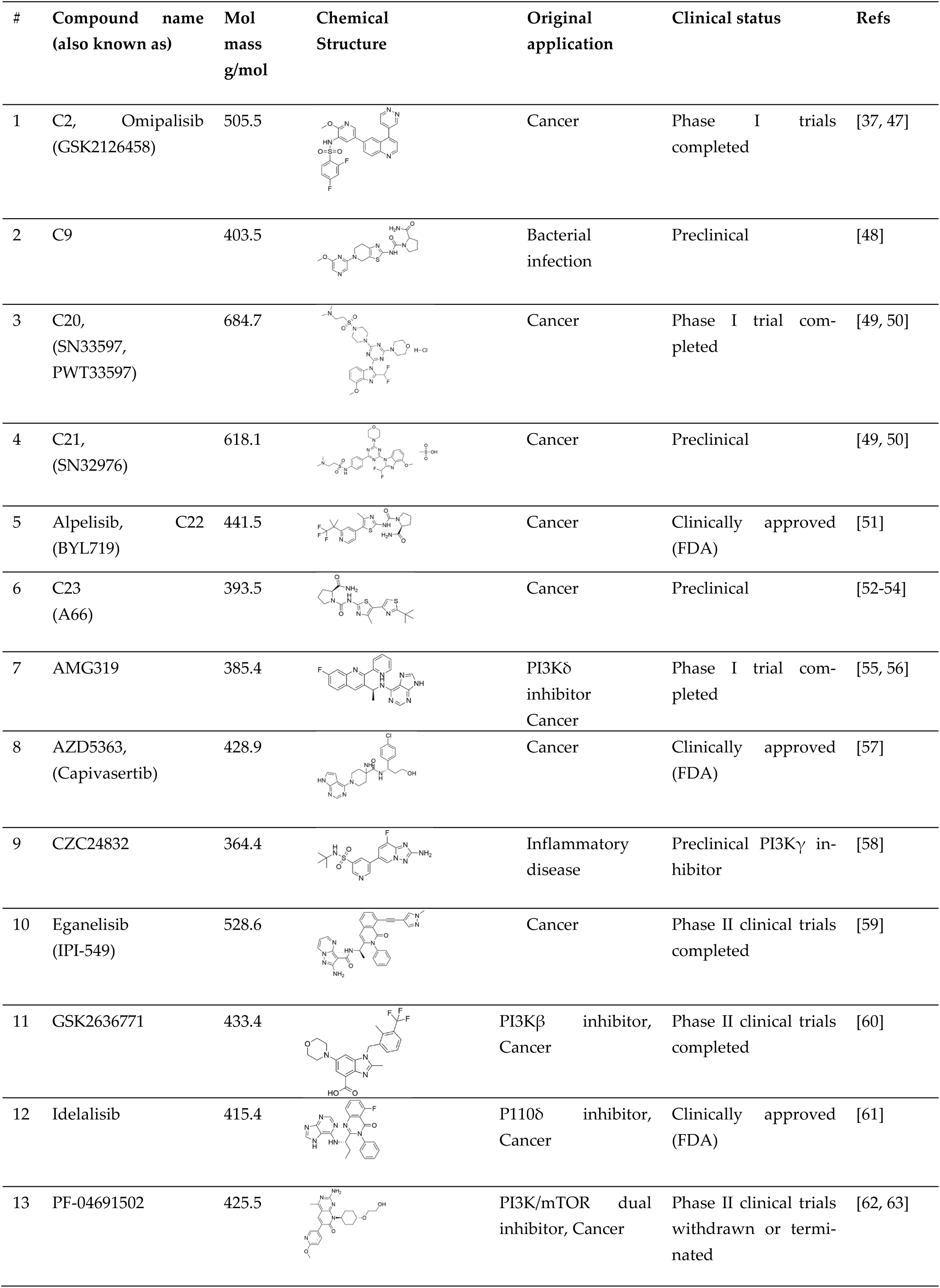
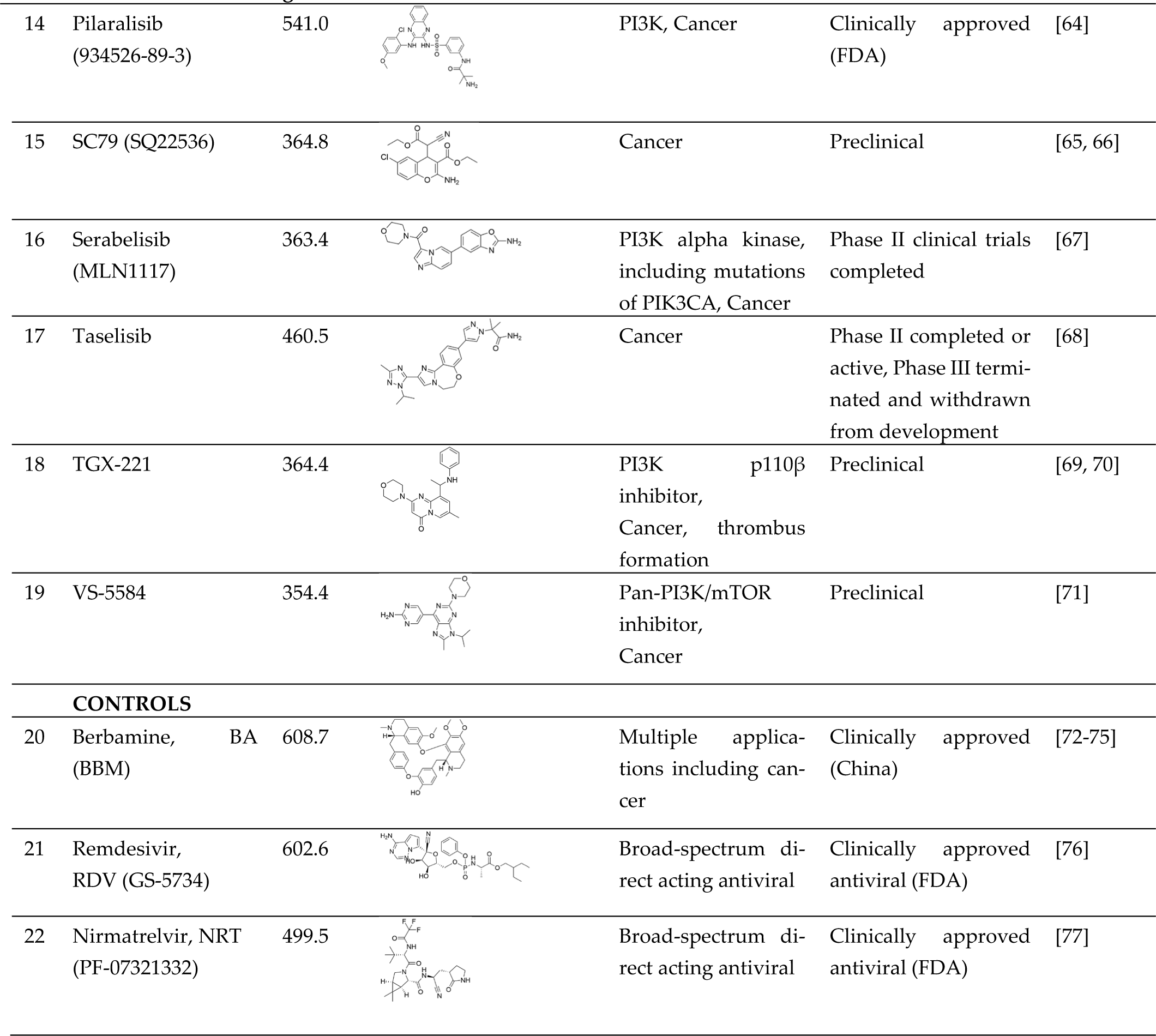
Compounds examined in this study.

### Cells, viruses, and approvals

The Murine 17Cl-1 cell line (derived from 3T3 cells, also known as NR-53719 *Mus musculus*), was supplied through kind support from BEI Resources. The MHV A59 strain and the NCTC clone 1469 cell line utilized for MHV cultivation were purchased from ATCC (Virginia, USA), VR-764. MRC5, Vero cells and human coronavirus OC43 (HCoV-OC43) were purchased from ATCC. SARS-CoV-2 variants Wuhan D614G, Delta NZ/21MV0256/2021 isolate, Mu NZ/CHCH/1673/2021 isolate and Omicron B.1.1.529 OM-1 were kind gifts from ESR New Zealand and University of Otago PC3 facility, New Zealand. All SARS-CoV-2 *in vitro* work was carried out under BSL3 conditions using biosafety approval BSC2847 for in the Faculty of Medical and Health Science’s PC3 facility at the University of Auckland.

### MHV culture assays

12-well plates were seeded with 6 × 10^4^ 17CL-1 cells and cultured overnight. The next day, cells were pre-treated with controls or test compounds at 0.5 μM for 4h then infected with MHV MOI 0.1. After 1h adsorption, virus was aspirated, cells washed and controls or test compounds at 0.5 μM were added and incubated for a further 14hpi. At 14hpi, the intracellular total RNA was extracted using the RNAfast200 Kit (Shanghai Feijie Co., Ltd, Cat#220011) as directed by the manufacturer, and RT-qPCR (Bio-Rad CFX96) was conducted to determine the viral RNA level, probing for the MHV non-structural protein 9 (nsp9). The negative controls of 0.1% v/v DMSO, and 10 μM BBM (Sigma-Aldrich) or 0.5 μM RDV (MedChemExpress, Monmouth Junction, NJ, USA) were used as positive controls.

### Viral plaque reduction assays

MRC5 (for HCoV-OC43 culture) or Vero (for SARS-CoV-2 culture) cells were seeded in 24 well plates and pre-treated with compounds or controls in Dulbecco’s modified Eagle’s medium (DMEM; Life Technologies, Carlsbad, CA, USA) with 2% (SARS-CoV-2) or 5% (HCoV-OC43) fetal bovine serum (FBS; Moregate Biotech, QLD, Australia), for 4h prior to infection unless stated otherwise. The compound vehicle DMSO 0.1% v/v was used as a negative inhibition (vehicle) control and positive inhibition controls included Berbamine (BBM) at 10-20 μM or Remdesivir (RDV) at 10 μM. After pre-treatment, cells were inoculated with HCoV-OC43 at MOI 0.03 or SARS-CoV-2 at MOI 0.0001, adsorbed for 1h then overlaid with media comprising test compounds or controls with 1.2% of sterile cellulose solution, 1x DMEM, 1x Glutamax (Thermo Fisher) and 1x non-essential amino acids (NEAA; Thermo Fisher Scientific, Waltham, MA, USA) for HCoV-OC43, 2% FBS (SARS-CoV-2) or 5% FBS (HCoV-OC43). After viral adsorption, virus inoculum was removed and 0.5 mL/well of cellulose-immobilising plaque overlay containing compounds or controls was added. Infected plates were incubated at 33°C, 5% CO_2_ for 120h (HCoV-OC43) or 37°C 5% CO_2_ for 72h (SARS-CoV-2). After incubation, cells were fixed with 10% formaldehyde and stained with 0.1% crystal violet in 20% methanol before enumeration of plaques. All graphs and IC_50_ values generated using Prism software v 10.3.1 (GraphPad, La Jolla, CA, USA).

### *In vitro* antiviral combination assays

SARS-CoV-2 plaque assays were completed as described above using the Omicron B.1.1.529 variant against a matrix of either RDV and/or C20 (1:1 ratio when combined) or NRT and/or C20 (1:1 ratio when combined) across a gradient of concentrations. The percentage of inhibition for each treatment as compared to the vehicle control (0.1% v/v DMSO) was used to calculate drug combinations using online software for interactive analysis and visualization of dose-response matrix data (http://www.synergyfinder.org/) using the Bliss model [45].

### Western blotting

Cells were lysed in cell lysis buffer with protease inhibitors, and protein concentration was measured by the Bradford Protein Assay. Proteins were loaded on SDS-PAGE gels, subjected to electrophoresis, and then transferred onto PVDF membranes (Millipore). Then, the membranes were blocked with 5% BSA in TBST (TBS buffer, containing 0.1% Tween-20) for 1h at room temperature. After incubating with indicated antibodies at 4°C overnight, the membranes were washed with TBST (3x5min). After incubating with appropriate amount of HRP-conjugated secondary antibodies for 1h at room temperature, images were captured by ChemiDoc using chemiluminescence substrates.

### Cytokine assays

Total RNA was extracted from tissues or cell lines by using the RNAfast200 Kit (Shanghai Feijie Co., Ltd, Cat#220011) as directed by the manufacturer, and quantitative real-time RT-PCR reactions were performed using the SYBR Premix Ex Taq Kit (Takara, Japan) with specific primers. Relative levels of each gene were normalized to that of 18s rRNA. The relative expressions of the target genes were calculated using the 2^−ΔΔCt^ method.

### Primary human tracheal epithelial cells (HTEC) culture

Human ethics approvals were through the Second Affiliated Hospital of Chongqing Medical University China. Human tracheal epithelial cells (HTECs) were isolated from patients undergoing bronchoscopy or surgical lung resection. Primary bronchial epithelial cultures were established using a well-established protocol [46]. Briefly, tissue fragments were dissociated via overnight digestion with 1.5 mg/mL pronase (Roche) at 4°C. The resulting cell suspension was centrifuged at 400 × g for 10 min at 4°C, and the pellet was resuspended in DNase solution for 5 min on ice, followed by another centrifugation step (400 × g, 10 min, 4°C). To remove contaminating fibroblasts, cells were resuspended in DMEM/F12 medium supplemented with 10% FBS and incubated at 37°C under 5% CO_2_ for 3-4 h. The supernatant containing epithelial cells was collected, resuspended in proliferation medium (*PneumaCult-Ex Plus*, STEMCELL Technologies, Canada), and seeded onto collagen-coated Transwell inserts. Proliferation medium was added to both apical and basal compartments. Upon reaching confluence, the medium was replaced with *PneumaCult-ALI* medium (STEMCELL Technologies) in the basal chamber only, creating an air-liquid interface (ALI) by exposing the apical surface. The monolayers were maintained at ALI for 20 additional days.

### Virus infection in HBEC

During viral challenge, cells were washed with culture medium to remove apical mucin. Subsequently, the virus was inoculated apically at a MOI of 2. Cells were incubated for 2h at 37°C to allow viral entry, after which the residual inoculum was removed by washing.

### Drug treatment

The basal chamber medium was replaced with drug solutions diluted in PneumaCult-ALI medium at the indicated concentrations for 2h. Following viral infection, the basal chamber medium was refreshed with drug-containing medium.

### Immunofluorescence staining

For immunofluorescence staining, plates containing Transwell® inserts were first immersed in ice-cold methanol (-20°C) and fixed overnight at -20°C. After fixation, inserts were blocked with 3% bovine serum albumin (BSA) in PBS for 1h at RT, then incubated with primary antibody for 2h at RT. Following three 5-minute PBS washes, samples were stained with fluorophore-conjugated secondary antibody for 1.5h at RT and washed again (3×5 min PBS). Nuclear counterstaining was performed using 4′,6-diamidino-2-phenylindole (DAPI) for 5 minutes at RT, followed by a final PBS wash. The liquid mountant was applied carefully to the membranes and coverslips were placed on top.

No generative artificial intelligence was used in this study or associated manuscript.

## 3. Results

### MHV RNA Levels Significantly Reduced by Dual Pan PI3K/m-TOR Inhibitors *In Vitro*

To establish a robust *in vitro* model for β-coronavirus infection, we first evaluated the replication kinetics of MHV-A59 in murine fibroblast 17CL-1 cells. Cells were infected with MHV-A59 at three different MOIs (0.01, 0.1, and 1), and viral titers were measured over time using a TCID₅₀ assay. As shown in the intercellular and extracellular growth curves (**Figure 1A**), MHV replicated efficiently in both compartments in a dose- and time-dependent manner. Intracellular viral titers peaked around 14 hpi, with the highest viral load observed at MOI 1. Extracellular viral release also increased steadily across all MOIs, reaching a maximum between 14 and 18 hpi. These findings confirm that MHV-A59 is highly infectious in 17CL-1 cells and capable of robust replication and viral shedding.

**Figure 1.**
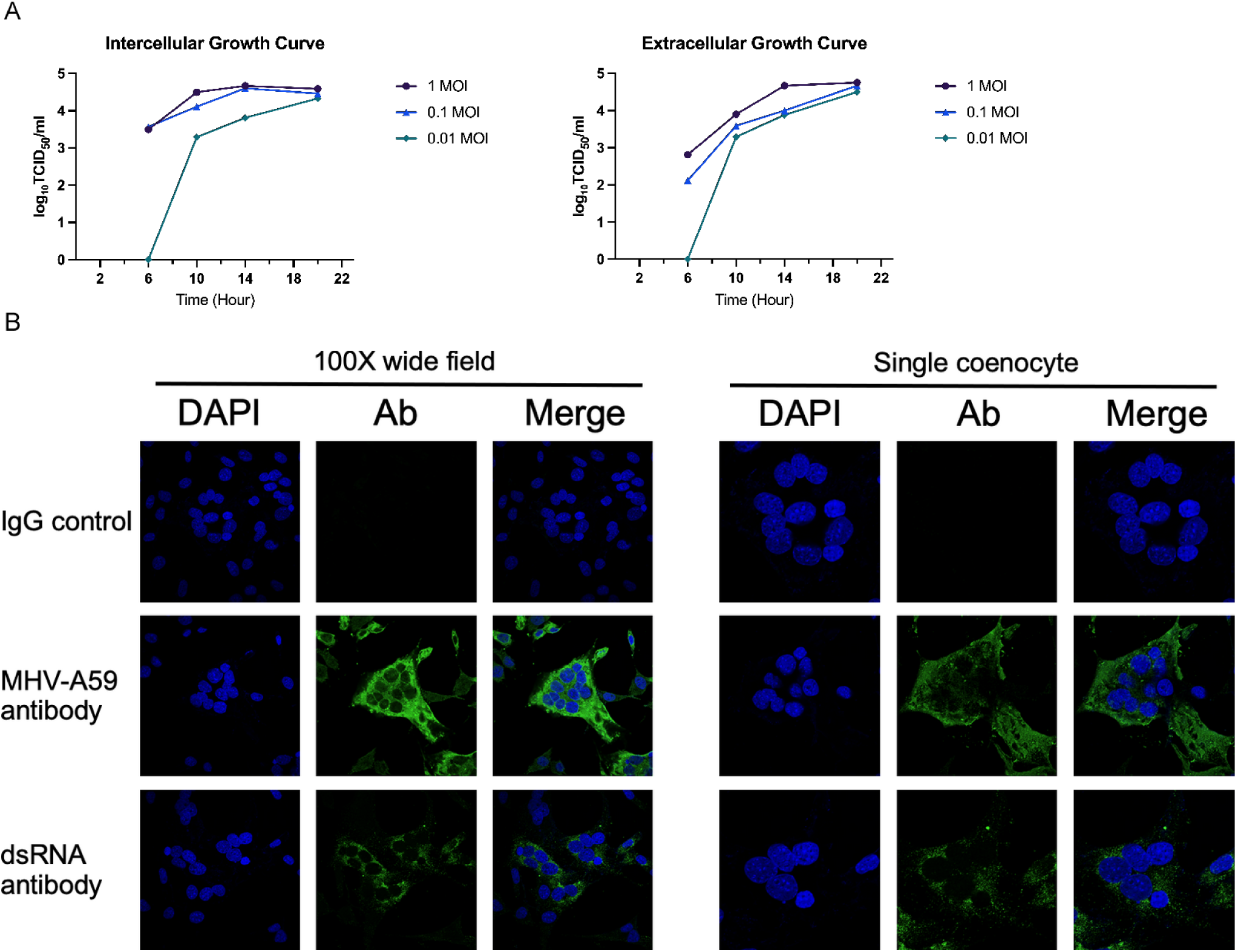
Mouse hepatitis virus (MHV), a β-coronavirus, infection model in host cells. (**A**) MHV-A59 efficiently infects 17CL-1 cells. (**B**) Immunofluorescence staining using antibodies against the MHV-N protein and double-stranded RNA (dsRNA) was used to assess the efficiency of MHV infection in host cells.

To visualize MHV infection at the cellular level, we performed immunofluorescence staining using antibodies against the MHV-A59 nucleocapsid (N) protein and double-stranded RNA (dsRNA), a marker of active viral replication. As shown in **Figure 1B**, both markers were strongly detected in infected cells, whereas no signal was observed in IgG isotype control samples. Wide-field (100×) and high-resolution single-cell images revealed the formation of multinucleated cells (coenocytes), a characteristic cytopathic effect of MHV infection. The clear green fluorescence co-localizing with nuclear DAPI staining in MHV-N and dsRNA-stained cells further confirmed productive viral infection. These results establish 17CL-1 cells as a suitable model for studying β-coronavirus infection dynamics and evaluating antiviral interventions.

To investigate the role of class I PI3K signaling in β-coronavirus infection, we first examined whether MHV-A59 activates this pathway in murine fibroblast 17CL-1 cells. Cells were infected with MHV (MOI of 0.1) and harvested at multiple time points for immunoblot analysis. As shown in **Figure 2A**, MHV infection rapidly induced phosphorylation of Akt at Ser473 and Thr308, as well as activation of downstream effectors such as 4EBP1, indicating robust activation of the PI3K-Akt-mTOR axis. Additionally, alterations in autophagy-related markers, including increased LC3-II and decreased p62, were observed over time, suggesting that viral infection also modulates autophagy. These findings support the hypothesis that MHV hijacks the PI3K signaling pathway to facilitate its replication and that targeting this pathway may offer therapeutic potential.

**Figure 2.**
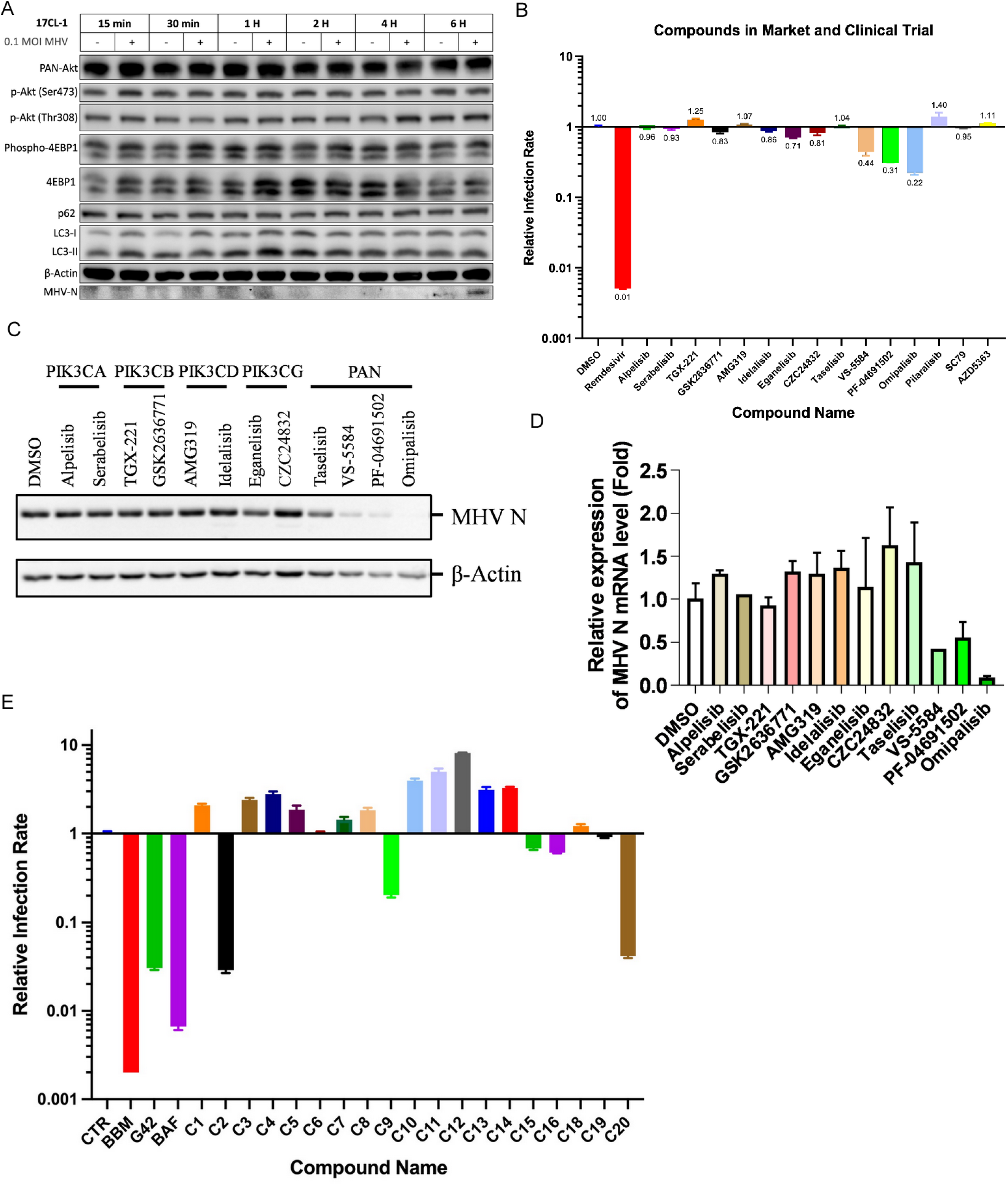
MHV infection activates the class I PI3K signaling pathway, and PI3K inhibitors suppress MHV replication. (A) Western blot analysis showing activation of PI3K-AKT-mTOR signaling and autophagy pathways in 17CL-1 cells infected with MHV-A59 (MOI = 0.1) at indicated time points (15 min to 6 h). Phosphorylation of AKT (Ser473 and Thr308), phospho-4EBP1, and changes in LC3-I/II and p62 levels indicate activation of PI3K signaling and modulation of autophagy. (B) Relative infection rates of MHV in 17CL-1 cells pretreated with 15 PI3K pathway inhibitors (0.5 μM), including isoform-specific and pan-PI3K inhibitors. MHV *nsp9* mRNA levels were quantified by RT-qPCR at 14 hpi and normalized to DMSO control. (C) Immunoblot analysis of MHV N protein expression following treatment with selected inhibitors from (B), representing different PI3K isoform classes. β-Actin was used as a loading control. (D) Quantification of MHV nucleoprotein mRNA levels by RT-qPCR after treatment with the inhibitors shown in (C), confirming antiviral effects at the transcriptional level. (E) Antiviral screening of an in-house library of 30 class I PI3K inhibitors. Compounds C2, C9, and C20 significantly reduced MHV infection at 0.5 μM, as measured by relative *nsp9* mRNA expression compared to control. Data in (B), (D), and (E) are shown as mean ± SD from at least three independent experiments.

To test this hypothesis, we conducted a targeted drug screen using a panel of 15 PI3K pathway inhibitors that are either FDA-approved or currently in clinical development. 17CL-1 cells were pretreated with each compound (0.5 μM) and then infected with MHV. Viral replication was assessed by quantifying *nsp9* mRNA levels relative to vehicle control (**Figure 2B**). Notably, the dual pan-PI3K/mTOR inhibitors **Omipalisib** (C2) and **PF-04691502** showed the strongest antiviral activity, reducing viral RNA levels by over 90%. Several isoform-specific inhibitors targeting PI3Kα, β, δ, and γ (e.g., Idelalisib, GSK2636771, Eganelisib) also exhibited moderate efficacy. These findings were confirmed at the protein level by immunoblotting for MHV N protein, which showed marked reduction in response to Omipalisib and PF-04691502 (**Figure 2C**). RT-qPCR analysis further validated that these compounds significantly suppressed viral gene expression (**Figure 2D**).

To expand the screening and identify additional candidates for preclinical development, we tested a custom in-house library of class I PI3K inhibitors (C1–C30) using the same infection model. As shown in **Figure 2E**, several compounds demonstrated potent antiviral activity at 0.5 μM, with **C2**, **C9**, and particularly **C20** standing out. C20 reduced MHV replication to 36% of the control, while C2 and C9 exhibited near-complete inhibition. These results not only validate dual inhibition of class I PI3K and mTOR as a promising host target for β-coronavirus intervention but also identify novel PI3K inhibitors with potential for repurposing or further optimization. Based on its strong antiviral effect and clinical development status, C20 was selected for detailed mechanistic and pharmacological evaluation in subsequent experiments.

To further characterize the antiviral potential and mechanism of action of the class I PI3K inhibitor **C20**, we first confirmed its ability to inhibit PI3K signaling in both murine macrophage RAW264.7 cells and fibroblast 17CL-1 cells. As shown in **Figures 3A and 3B**, C20 treatment markedly reduced the phosphorylation of Akt at Ser473 and Thr308, as well as phosphorylation of the downstream effector 4EBP1, in a time-dependent manner. These findings validate C20 as a potent inhibitor of class I PI3K-Akt-mTOR signaling in multiple MHV-permissive cell types, establishing a foundation for evaluating its antiviral effects.

**Figure 3.**
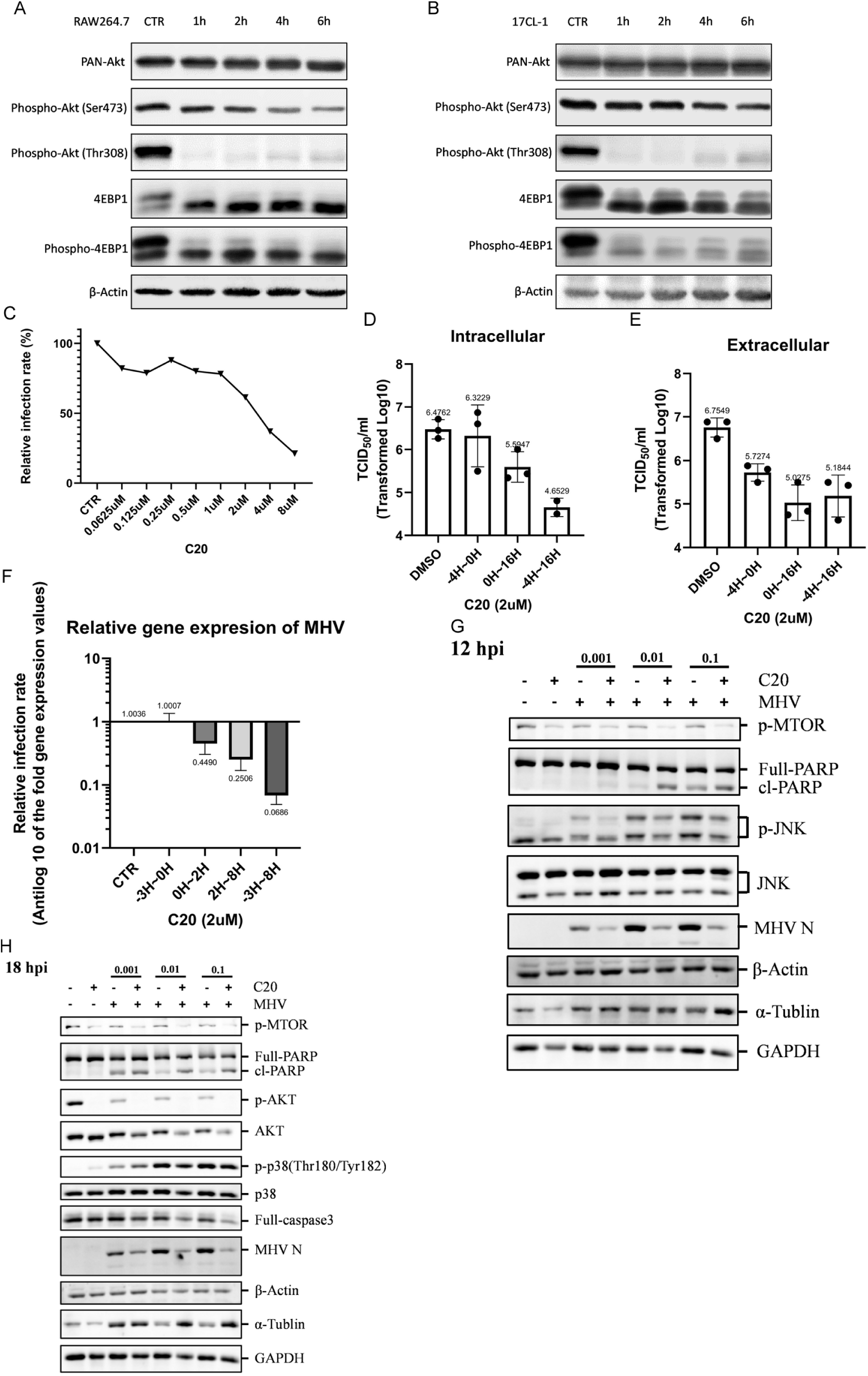
The class I PI3K inhibitor C20 suppresses MHV replication in a dose- and time-dependent manner and modulates host signaling pathways. (A–B) Western blot analysis showing inhibition of PI3K signaling in RAW264.7 cells (A) and 17CL-1 cells (B) treated with 2 μM C20 for the indicated time points (1–6 h). C20 reduced phosphorylation of Akt (Ser473 and Thr308) and 4EBP1, confirming effective inhibition of the PI3K-Akt-mTOR axis. (C) Dose–response curve showing relative infection rate (%) of MHV in 17CL-1 cells treated with increasing concentrations of C20, as determined by *nsp9* mRNA expression. (D–E) TCID₅₀ assays showing intracellular (D) and extracellular (E) viral titers in 17CL-1 cells pretreated with DMSO or C20 (2 μM), added at 0 h or 4 h post-infection (hpi) and harvested at 18 hpi. (F) Time-of-addition analysis of C20 (2 μM), demonstrating significant inhibition of MHV *nsp9* gene expression when added up to 8 hpi. (G–H) Western blot analysis at 12 hpi (G) and 18 hpi (H) showing that C20 treatment reduces phosphorylation of mTOR, JNK, and p38, as well as expression of MHV N protein. C20 also decreases levels of cleaved PARP and cleaved caspase-3, suggesting modulation of apoptosis. β-Actin, α-Tubulin, and GAPDH were used as loading controls.

To determine the dose-response relationship of C20 against MHV infection, 17CL-1 cells were treated with increasing concentrations of C20 prior to viral infection. As shown in **Figure 3C**, C20 inhibited MHV replication in a dose-dependent manner, with significant suppression observed at concentrations

≥1 μM. Viral titers measured at 18 hpi further confirmed C20’s inhibitory effect in both intracellular (**Figure 3D**) and extracellular (**Figure 3E**) compartments. Time-of-addition experiments demonstrated that administering C20 up to 8 hours post-infection still significantly reduced MHV mRNA levels (**Figure 3F**), suggesting that C20 functions not only as a viral entry blocker but also interferes with postentry replication stages. Immunoblot analyses at 12 and 18 hpi further revealed that C20 suppressed MHV N protein expression and reduced phosphorylation of mTOR, JNK, and p38 MAPK (**Figures 3G and 3H**), while also modulating apoptotic markers such as cleaved PARP and cleaved caspase-3. These data indicate that C20 likely exerts antiviral effects by disrupting both viral replication machinery and host stress response pathways.

In summary, these results demonstrate that C20 potently inhibits MHV replication through direct suppression of class I PI3K signaling and downstream pro-viral pathways, including mTOR and MAPK cascades. The ability of C20 to retain antiviral activity even when added several hours after infection highlights its therapeutic potential as a post-exposure antiviral agent. Furthermore, its modulation of host cell apoptosis and inflammation pathways suggests that C20 may provide dual benefits: controlling viral load and limiting virus-induced cytopathology.

### Selected PI3K Inhibitors Show Antiviral Activity Against Human Coronavirus OC43 *In Vitro*

To determine whether class I PI3K inhibitors identified in the MHV screening also exhibit antiviral activity against human coronaviruses, we evaluated the effect of selected compounds on **HCoV-OC43** infection *in vitro*. As shown in **Figure 4A**, a plaque reduction assay revealed that several compounds, particularly **C20**, markedly suppressed HuCoV-OC43 plaque formation in a dose-dependent manner. These results indicate that the antiviral activity of the PI3K inhibitor C20 is not limited to MHV, but extends to clinically relevant human β-coronaviruses.

**Figure 4.**
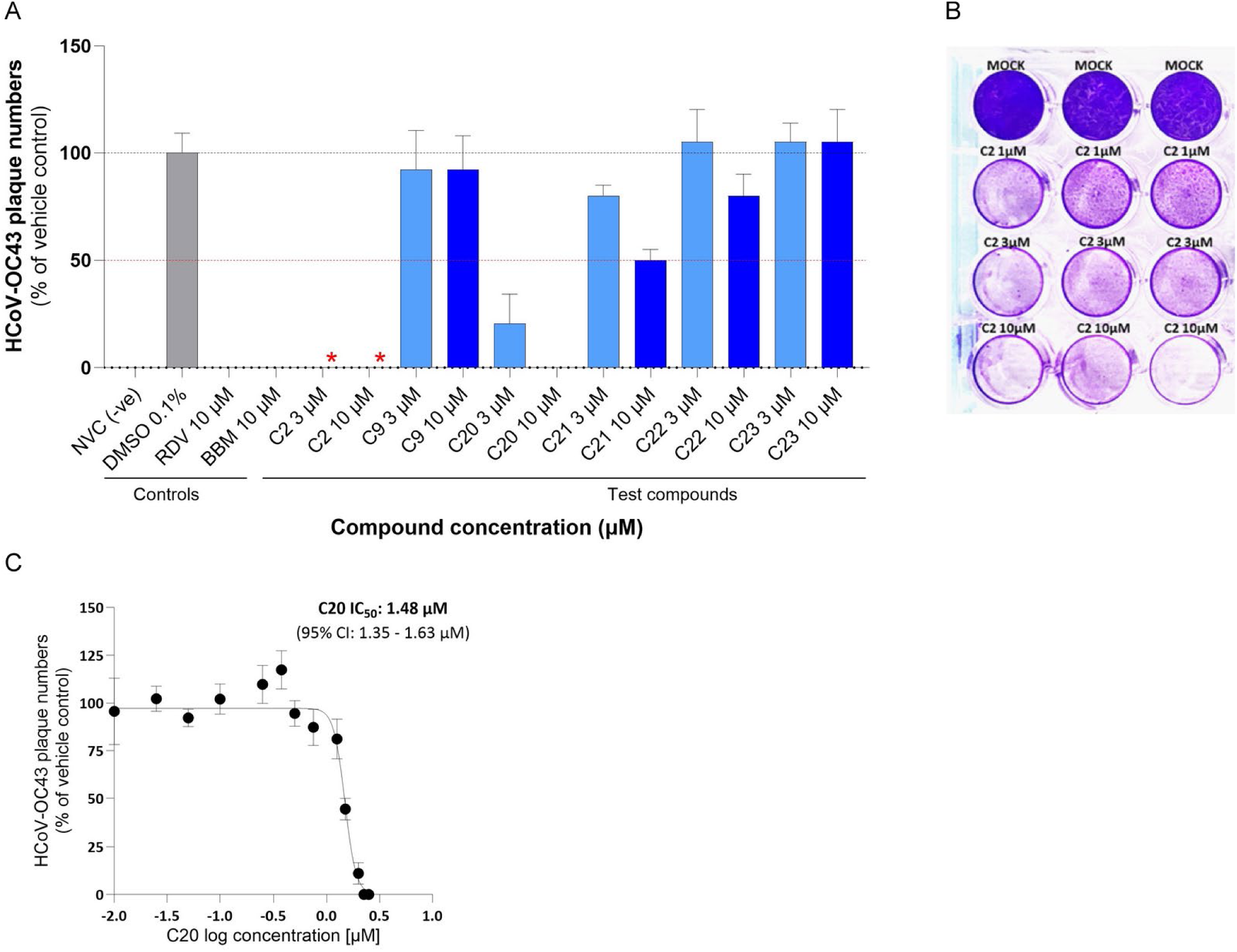
PI3K inhibitor C20 exhibits potent antiviral activity against human coronavirus OC43. (A) HCoV-OC43 plaque reduction assay results. OC43-infected cells were treated with selected test compounds at indicated concentrations. Plaque numbers were normalized to DMSO (0.1%) vehicle control. Positive controls included remdesivir (RDV, 10 μM) and berbamine (BBM, 10 μM) [73, 75]. C20 significantly reduced plaque numbers in a dose-dependent manner, while C2 was cytotoxic to the MRC5 cells (red asterisks). (B) Representative crystal violet-stained wells showing cytopathic effects of C2 (Omipalisib) on MRC5 cells with treatment, indicating the lack of a selective index (1, 3, and 10 μM). (C) Dose–response curve for C20 against HCoV-OC43. Plaque numbers were quantified and plotted as a percentage of vehicle control. The calculated EC₅₀ value for C20 was 1.48 μM (95% CI: 1.35–1.63 μM). The mean of triplicate datasets from three independent experiments are shown ±SD.

Despite the antiviral activity of C2 against MHV, when examined against HuCoV-OC43 at 3 and 10 μM, C2 demonstrated significant cell thinning, making plaques uncountable. Visual confirmation of cytotoxic thinning of the cells was demonstrated in **Figure 4B**, where crystal violet staining of infected monolayers showed substantial toxicity by C2 at 3 μM and 10 μM compared to untreated or mock-infected controls. As C2 did not demonstrate any selective index, it was dropped from further evaluation against HuCoV-OC43.

To further quantify the antiviral potency of the lead compound C20 against HCoV-OC43, we performed a dose–response analysis to determine its EC₅₀. As shown in **Figure 4C**, C20 displayed a sigmoidal dose–response curve with an **EC₅₀ of 1.48 μM** (95% CI: 1.35-1.63 μM), confirming its strong inhibitory effect at low micromolar concentrations. Notably, at higher concentrations, cell morphology appeared compromised, suggesting cytotoxicity beyond effective antiviral doses. Together, these data demonstrate that C20 is a potent inhibitor of OC43 infection, supporting its potential as a **broad-spectrum antiviral candidate** targeting class I PI3K-dependent and mTOR-dependent viral replication mechanisms.

### C20 has Antiviral Activity against Four SARS-CoV-2 Variants *In Vitro*

We screened a small panel of PI3K-targeting compounds using a plaque reduction assay against the Wuhan D614G variant in Vero cells. As shown in **Figure 5A**, both C2 (Omipalisib) and C20 demonstrated inhibitory effects against viral replication, with C2 completely blocking plaque formation at concentrations as low as 1 μM, though it was associated with notable thinning of the cell monolayer, indicating potential cytotoxicity and a low selective index. In contrast, C20 exhibited a more gradual and concentration-dependent antiviral effect, reducing plaque formation by 33.3–45.8% at 1–3 μM and completely abolishing detectable plaques at 10 μM without severe cytopathic effects. To quantify this effect, a dose–response curve established an IC₅₀ of **5.94 μM** (95% CI: 2.86–7.44 μM) for C20 against the Wuhan variant (**Figure 5B**).

**Figure 5.**
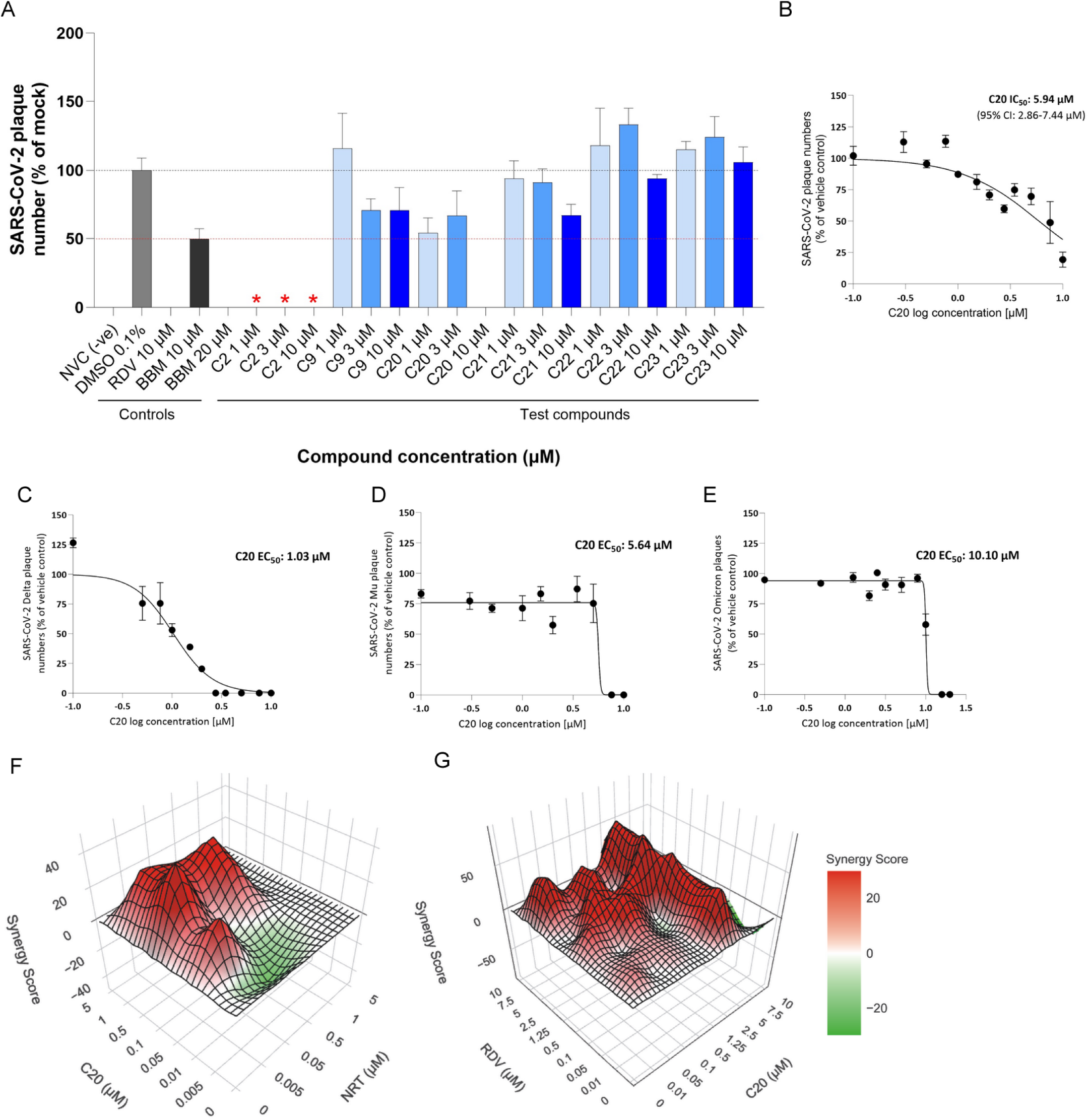
C20 exhibits broad-spectrum antiviral activity against SARS-CoV-2 variants and synergizes with direct-acting antivirals. (A) SARS-CoV-2 (Wuhan D614G) plaque reduction assay in Vero cells treated with PI3K inhibitors (1, 3, or 10 μM). Plaque counts are shown as a percentage of vehicle (DMSO 0.1%) control. Positive controls include Remdesivir (RDV, 10 μM) and Berbamine (BBM, 10 μM). Red asterisks indicate cell monolayer thinning due to cytotoxic effects of the drug. (B–E) Dose–response curves of C20 against (B) Wuhan D614G (IC₅₀ = 5.94 μM), (C) Delta variant (IC₅₀ = 1.03 μM), (D) Mu variant (IC₅₀ = 5.64 μM), and (E) Omicron variant (IC₅₀ = 10.10 μM). Data represent mean of triplicate datasets from at least two independent experiments ± SD. (F–G) Bliss synergy analysis of C20 in combination with (F) NRT or (G) RDV in SARS-CoV-2 (Wuhan) plaque assays. 3D synergy plots show positive Bliss scores (red) indicating synergistic antiviral effects. Mean Bliss scores were 18 (C20 + NRT) and 6 (C20 + RDV).

To assess whether C20’s antiviral activity extended across SARS-CoV-2 variants, we tested it against three additional strains: Delta (NZ/21MV0256/2021), Mu (NZ/CHCH/1673/2021), and Omicron (B.1.1.529). C20 showed effective inhibition across all variants with IC₅₀ values of **1.03 μM**, **5.64 μM**, and **10.10 μM**, respectively (**Figures 5C–5E**). These results confirm that C20 retains broad-spectrum antiviral activity across genetically and phenotypically distinct SARS-CoV-2 lineages, particularly those of public health concern.

Given its promising activity, we next explored whether C20 synergizes with existing FDA-approved direct-acting antivirals. Using a Bliss independence model, we tested C20 in combinatorial matrices with **Nirmatrelvir (NRT)**, a viral protease inhibitor, and **Remdesivir (RDV)**, a viral polymerase inhibitor. As shown in the 3D synergy plots (**Figures 5F and 5G**), C20 exhibited overall **synergistic effects** when combined with both NRT (mean Bliss score = 18) and RDV (mean Bliss score = 6). While some antagonistic effects were observed at lower concentrations, the net interactions were positive, highlighting C20’s potential as part of a combination therapy to enhance treatment efficacy and reduce resistance risks. These findings position C20 as a promising host-targeting broad-spectrum antiviral with synergistic potential alongside frontline SARS-CoV-2 therapies.

### Evaluating the Antiviral Activity of C20 Against SARS-CoV-2 Using a Human Lung Organoid Model

To model the primary targets of Omicron infection in the human airway, we established air–liquid interface (ALI) cultures of primary human tracheal epithelial cells (HTECs). These cultures differentiate into ciliated, goblet, and basal cells that mimic the native architecture of the upper respiratory epithelium. Immunofluorescence staining confirmed that the ALI-cultured HTEs express key markers associated with SARS-CoV-2 susceptibility [46]. As shown in the left panel (Figure S1), cells stained positively for acetylated tubulin (AC-tubulin), a marker of differentiated ciliated epithelial cells, indicating successful formation of physiologically relevant airway epithelium. In the right panel (Figure S1), the same cultures showed strong surface expression of ACE2, the primary host receptor for SARS-CoV-2, confirming that these organoid models are well-suited for studying infection and drug response.

**Figure S1.**
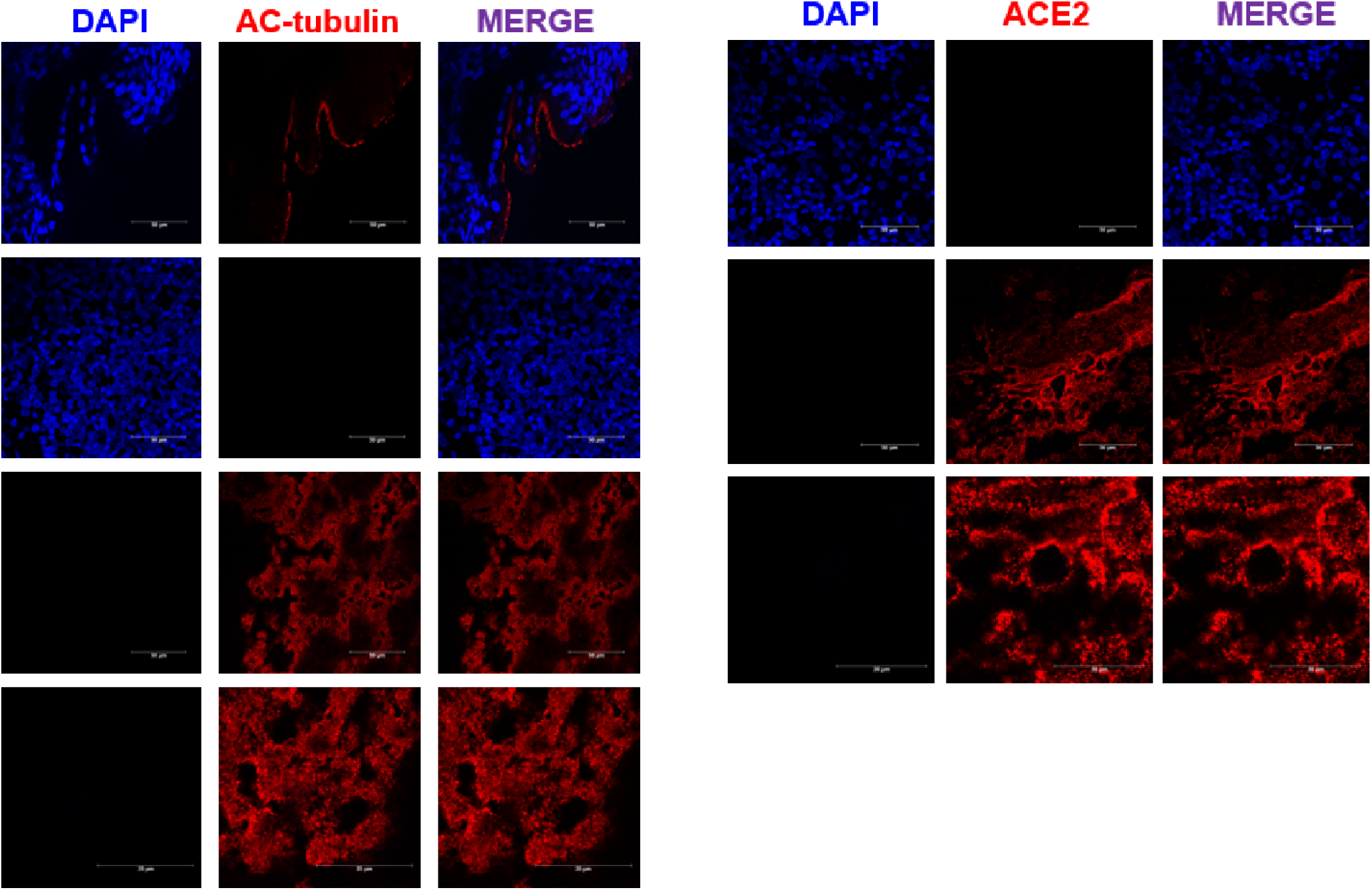
ALI-cultured human tracheal epithelial cells express cilia marker AC-tubulin and SARS-CoV-2 receptor ACE2. Representative immunofluorescence images of HTECs grown at an air–liquid interface (ALI), showing expression of (left) acetylated tubulin (AC-tubulin, red) and (right) ACE2 (red). Nuclei were counterstained with DAPI (blue). Merged images confirm the presence of ciliated cells and ACE2-expressing surfaces in differentiated HTECs. These results validate the use of ALI-cultured HTECs as a physiologically relevant model for studying SARS-CoV-2 infection. Scale bars = 50 μm.

These findings support the rationale for using ALI-cultured HTECs to evaluate the infectivity of SARS-CoV-2 variants such as Wuhan and Omicron, which predominantly target ciliated cells of the upper airway. The robust co-expression of AC-tubulin and ACE2 in these cells suggests they are highly permissive to viral entry and replication. In subsequent experiments (described in Figure 6), infection by both Wuhan and Omicron variants confirmed their tropism for AC-tubulin+ cells. Moreover, antiviral testing demonstrated that treatment with C20, similar to remdesivir, effectively suppressed infection in these ALI-cultured systems. Together, these results validate the use of ALI-HNE/HTE organoids as a physiologically relevant platform for assessing SARS-CoV-2 variant infectivity and host-targeted antiviral efficacy.

**Figure 6.**
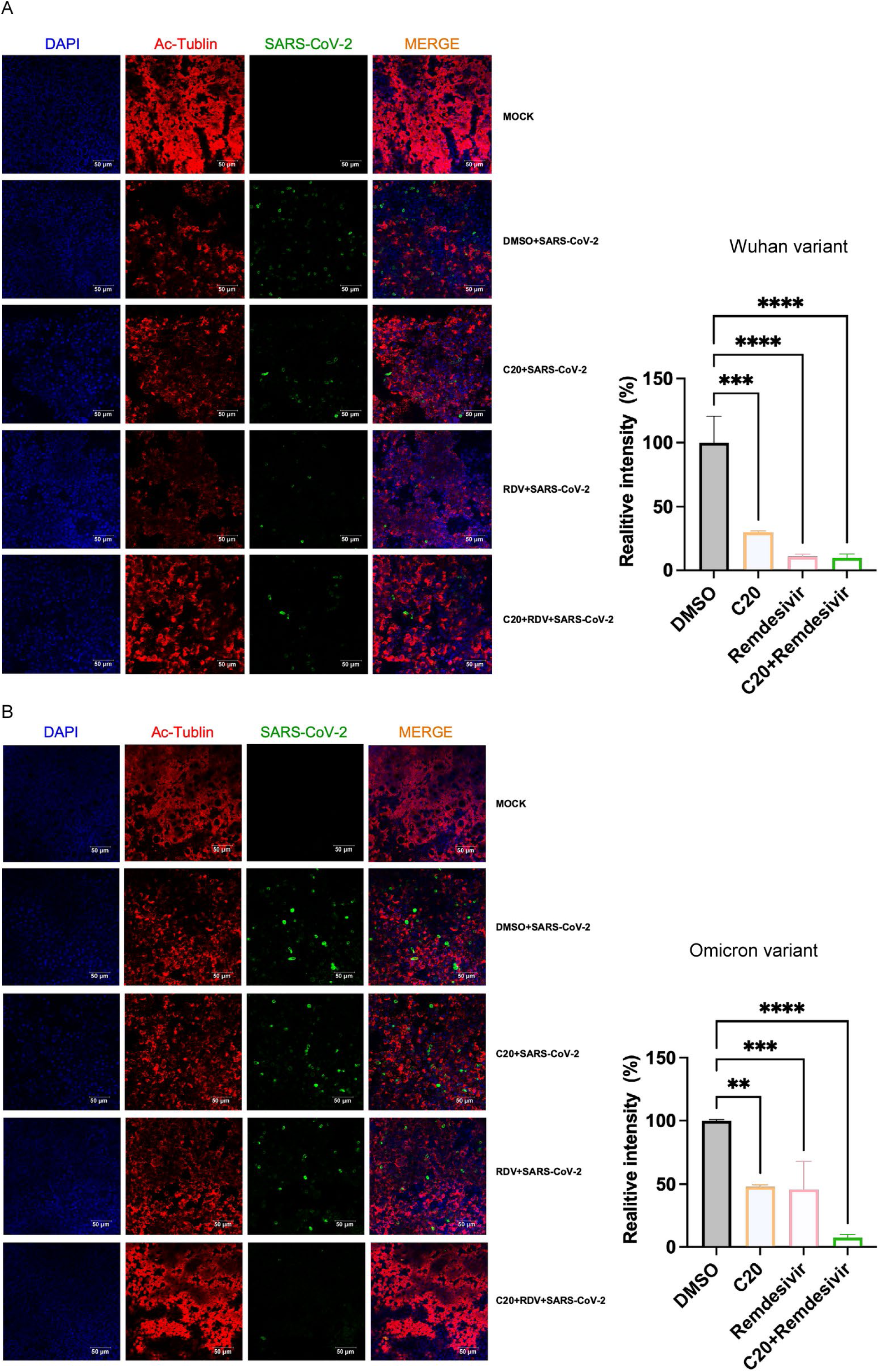
C20 inhibits SARS-CoV-2 Wuhan and Omicron variant infection in ALI-cultured human tracheal epithelial cells. (A–B) Immunofluorescence staining of air–liquid interface (ALI)–cultured human tracheal epithelial cells (HTECs) infected with either (A) SARS-CoV-2 Wuhan variant or (B) Omicron variant for 24 hours. Cells were treated with DMSO, C20 (1.0 μM), remdesivir (0.5 μM), or their combination. Samples were stained for acetylated tubulin (Ac-tubulin, red), SARS-CoV-2 N protein (green), and nuclei (DAPI, blue). Merged images show colocalization of viral antigen with ciliated epithelial cells. Bar graphs on the right show quantification of SARS-CoV-2 signal intensity relative to DMSO-treated controls. Both C20 and remdesivir significantly reduced viral infection, and the combination treatment achieved near-complete suppression. Data represent mean ± SD from three independent experiments; p < 0.01, *p < 0.001, **p < 0.0001 by one-way ANOVA. Scale bars = 50 μm.

To evaluate the antiviral efficacy of C20 against SARS-CoV-2 in a physiologically relevant model, we used ALI-cultured human tracheal epithelial cells (HTEs) and assessed infection by the Wuhan (**Figure 6A**) and Omicron (**Figure 6B**) variants via immunofluorescence staining. In DMSO-treated infected controls, robust SARS-CoV-2 N protein staining (green) was observed, predominantly colocalized with the cilia marker acetylated tubulin (red), indicating successful viral targeting of ciliated epithelial cells. Treatment with C20 or remdesivir significantly reduced viral antigen levels, while the combination of C20 and remdesivir led to near-complete suppression of viral signal. Quantification of fluorescence intensity confirmed a marked reduction in viral load with either treatment alone and an even greater effect when combined, suggesting additive or synergistic antiviral activity.

These findings support that C20 effectively inhibits SARS-CoV-2 infection in ALI-cultured airway epithelium, including for both the ancestral Wuhan strain and the highly transmissible Omicron variant. The reduction in viral antigen levels in the C20 and remdesivir co-treatment group was statistically significant compared to monotherapies, particularly in the Omicron setting, where viral resistance to some therapeutics is a concern. These results not only highlight the antiviral potency of C20 in a physiologically relevant model but also suggest that combining host-targeting agents like C20 with direct-acting antivirals such as remdesivir may enhance therapeutic efficacy against current and emerging SARS-CoV-2 variants.

### C20 Enhances Cytokine Expression in MHV-Infected Cells

To investigate how MHV infection influences host cytokine expression and whether the PI3K inhibitor C20 modulates this response, we first assessed time-dependent changes in cytokine gene expression in **17CL-1 fibroblast cells**, a permissive cell line for MHV-A59 infection. As shown in **Figure 7A**, MHV RNA levels progressively increased from 16 to 24 hpi, confirming productive viral replication. Alongside viral amplification, we observed a time-dependent increase in several proinflammatory and antiviral cytokines, including IL-1β, IL-6, IL-10, IFN-γ, and to a lesser extent IFN-β and TNF-α. These results suggest that MHV infection in epithelial-like cells initiates a delayed but coordinated cytokine response that could contribute to antiviral signaling or inflammation.

**Figure 7.**
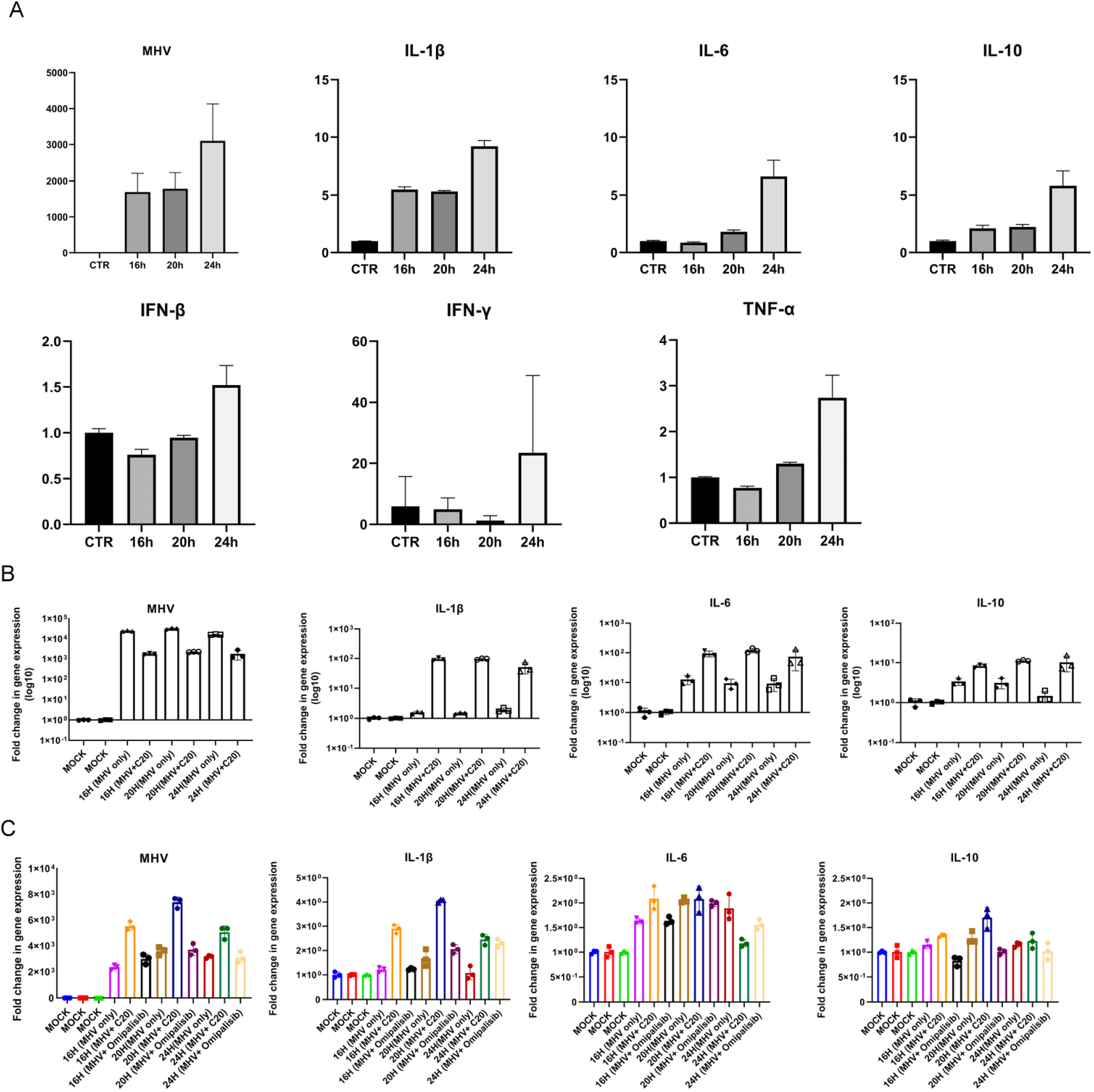
C20 enhances cytokine expression in MHV-infected 17CL-1 fibroblasts and RAW264.7 macrophages. (A) Time-course analysis of MHV replication and cytokine gene expression in 17CL-1 fibroblast cells infected with MHV-A59 (MOI = 0.1). Cells were harvested at 16, 20, and 24 hours post-infection (hpi), and mRNA levels of MHV, IL-1β, IL-6, IL-10, IFN-β, IFN-γ, and TNF-α were quantified by RT-qPCR. Results show progressive viral replication and a delayed increase in inflammatory cytokine expression. (B) Gene expression in RAW264.7 macrophages infected with MHV (MOI = 0.1) for 24h, with or without C20 (2 μM) pretreatment. C20 enhanced the expression of IL-1β, IL-6, and IL-10 in infected cells without significantly reducing viral RNA levels. (C) Expanded validation of cytokine responses in RAW264.7 cells across multiple treatment conditions, including MHV alone, C20 alone, and C20 + MHV. C20 significantly increased cytokine gene expression when combined with viral infection. Data are presented as mean ± SD from at least three independent experiments.

We next examined the effect of C20 on cytokine responses in **RAW264.7 macrophages**, a cell type that plays a central role in innate immunity. As shown in **Figure 7B**, MHV infection alone induced a strong upregulation of MHV RNA and cytokines such as IL-1β, IL-6, and IL-10. Notably, pretreatment with C20 (2 μM) further amplified cytokine expression in MHV-infected macrophages, without significantly affecting viral RNA levels. This enhancement of cytokine response was validated in an expanded panel of conditions in **Figure 7C**, where the combination of C20 and MHV consistently led to elevated expression of IL-1β, IL-6, and IL-10 compared to MHV alone or C20 alone.

In summary, these findings demonstrate that C20 does not impair MHV replication in epithelial or macrophage cell lines, but rather **augments host cytokine responses**, particularly in immune cells. This suggests that C20 may exert dual activity: acting as a PI3K inhibitor with antiviral potential, while simultaneously enhancing innate immune activation. Such immunomodulatory effects may be beneficial in the context of viral infection, although further studies are required to balance these benefits with the risk of inflammatory overactivation.

## 4. Discussion

The emergence of SARS-CoV-2 variants with partial resistance to current therapeutics such as molnupiravir, remdesivir, and nirmatrelvir underscores the urgent need for new antiviral strategies. Host-targeting antivirals, such as PI3K inhibitors, offer a promising path forward by interfering with conserved cellular pathways exploited by diverse viruses. Moreover, combining such host-targeting agents with direct-acting antivirals could reduce required doses, mitigate side effects, and importantly, delay or prevent the emergence of viral resistance.

Here we investigated the antiviral effects of a range of classes of PI3-kinase inhibitors against coronaviruses. None of the isoform selective class I PI3-kinase inhibitors had antiviral activity. However, we demonstrate dual class I PI3K and mTOR inhibitor C20 has both an antiviral and immunomodulatory agent against coronaviruses, including SARS-CoV-2 and MHV. We demonstrated that C20 exhibits potent antiviral activity across multiple SARS-CoV-2 variants—including Wuhan, Delta, Mu, and Omicron—and that it retains efficacy in both transformed cell lines and physiologically relevant air–liquid interface (ALI) airway models. Notably this phenocopies results observed with omipalisib, another PI3K/mTOR inhibitor [37, 39].

One concern is whether there is a feasible therapeutic window for PI3K/mTOR inhibitors *in vivo*. While there are no published reports of the toxicity profile of C20, it has been through Phase I trials and used in preclinical murine models indicating there are safe levels for dosing. However, C20’s cytostatic mechanism and low micromolar IC₅₀ and CC₅₀ values suggests possible toxicity when used at high doses, particularly in vulnerable patients with SARS-CoV-2. However, any such toxicities could be reduced by use in synergistic combination with other antivirals. Indeed, C20 showed synergistic antiviral effects when combined with FDA-approved antivirals such as remdesivir and nirmatrelvir, supporting its potential as a combinatorial therapeutic option, that may allow for the lowering the dose of C20 needed and thus avoiding potential toxicities.

Another interesting and important observation of the current study relates to the impacts of C20 in modulating the host immune response, particularly cytokine production. Our results in RAW264.7 macrophages and 17CL-1 fibroblasts showed that C20, while not suppressing viral RNA levels in all cases, significantly enhanced expression of cytokines such as IL-1β, IL-6, and IL-10 in MHV-infected cells. These data may initially appear contradictory in the context of cytokine storm—one of the major contributors to severe SARS-CoV-2 pathology. However, this effect could also be viewed as beneficial, especially in the early stages of infection where robust innate immune activation is crucial for viral control. This immunostimulatory property of C20 may enhance early antiviral responses, serving as a tool to limit viral replication and spread before adaptive immunity is fully established.

That said, the immunomodulatory effects of PI3K inhibitors must be carefully contextualized. While compounds like Duvelisib are being tested to suppress immune-driven lung injury in SARS-CoV-2 infection [78], others such as C20 may amplify cytokine production and potentially exacerbate inflammatory damage in later stages. Thus, their application may depend heavily on timing, dosage, and patient stratification. It is also worth noting that several other PI3K/AKT/mTOR inhibitors—such as Omipalisib, Capivasertib, and Dactolisib—have demonstrated *in vitro* efficacy against SARS-CoV-2, and impacts on the host immune system have been proposed but not yet confirmed [79-81]. The clinical development of these PI3K inhibitors as antivirals has largely stalled due to either cytotoxicity, limited *in vivo* efficacy, or off-target effects such as cardiac toxicity or immunosuppression.

In summary, C20 shows potential as a broad-spectrum antiviral agent with both direct-acting and immunomodulatory properties. Its ability to synergize with existing clinical antivirals, coupled with its favorable tolerability profile, warrants further investigation in preclinical and clinical models. Given the complex role of PI3K signaling in both viral replication and immune regulation, future studies should focus on delineating the optimal therapeutic window and understanding its effects in different disease stages to fully realize its clinical potential.

## Summary

In this study, we evaluated a panel of PI3K inhibitors for their broad-spectrum antiviral potential against several coronaviruses with distinct tropisms, including HCoV-OC43, SARS-CoV-2, and MHV. Among these, C20 emerged as a leading candidate, demonstrating consistent antiviral activity across all coronaviruses examined, with IC₅₀ values in the low micromolar range. Notably, C20 inhibited replication of four major SARS-CoV-2 variants—Wuhan, Delta, Mu, and Omicron—supporting its potential as a variant-resilient SARS-CoV-2 therapeutic.

Beyond its direct antiviral effects, C20 also enhanced cytokine expression in virus-infected cells, suggesting a dual role as both an antiviral and immunomodulatory agent. The ability of C20 to suppress infection in physiologically relevant airway organoid models and synergize with approved antivirals remdesivir and nirmatrelvir further strengthens its translational value. Taken together, these findings warrant further preclinical and clinical investigation of C20 as a promising broad-spectrum, host-targeted antiviral therapy.

## Supplementary Materials

Figures S1 and S2

## Author Contributions

JY, NN, QW and PS contributed to the conception of the study; JX, NC, BSH, JT, AN and NN contributed to data acquisition; JX, NC and NN analyzed the data; NN and JY wrote the manuscript; GN and JF synthesized and provided the small molecule compounds; QW, JF and PS contributed to review and editing the manuscript. All authors provided comments on manuscript drafts and approved the final version.

## Data Availability Statement

Data available on request.

## Acknowledgments

We acknowledge the support of the University of Auckland PC3 Facility and associated staff including Dr Shabihah Shahrudin, Rebecca Marnane, Dr Claire Honney and Benedict Uy. We also acknowledge BEI Resources (NIAID, NIH), a component of the Human Microbiome Project, for providing the Murine 17Cl-1 cell line.

